# The biomolecular profiles of extracellular vesicles from odontogenic stem cell lines depict donor-dependent differences and emphasize their therapeutic and regenerative potential

**DOI:** 10.1101/2025.05.13.653575

**Authors:** Yong Xu, Alp Sarisoy, Eva Miriam Buhl, Amanda Salviano-Silva, Julian Gonzalez Rubio, Anja Lena Thiebes, Sandra Fuest, Ralf Smeets, Franz Lennard Ricklefs, Christian Apel

**Author notes:** Correspondence: Christian Apel, Department of Biohybrid & Medical Textiles (BioTex), Center for Biohybrid Medical Systems (CBMS), Institute for Applied Medical Engineering, RWTH Aachen University, Forckenbeckstr. 55, 52074, Aachen, Germany.

## Abstract

**Background:** Stem cell-derived extracellular vesicles (EVs) hold great promise in regenerative medicine. However, a comprehensive understanding of the regenerative capabilities of EVs from different stem cell sources remains limited.

**Methods:** This study systematically compares EVs derived from three odontogenic cell types. Analyses includes EV isolation and characterization, cell viability assays, vasculogenesis experiments, proteomic profiling, and miRNA sequencing.

**Results:** All three EV types displayed similar surface marker profiles. Dental pulp stem cell-derived EVs showed superior cellular uptake, promoted higher cell proliferation, and enhanced vasculogenesis compared to periodontal ligament stem cell-derived EVs. Gingival fibroblast-derived EVs performed similarly in functional assays. Principal component analysis of miRNA profiles revealed strong biological heterogeneity among EV sources, with donor-specific factors exerting a greater influence on EV characteristics than cellular origin—an aspect underexplored in prior studies.

**Conclusions:** These findings underscore the complexity of EV functionality and highlight the regenerative potential of dental stem cell-derived EVs.

## Introduction

Research on extracellular vesicles (EVs) has advanced significantly, revealing their potential as therapeutic targets, biomarkers, innovative drug delivery systems, and stand-alone therapeutic agents. Mesenchymal stem cells (MSC) are currently the primary source of native EVs for clinical applications in tissue engineering and regenerative medicine. MSC-derived EVs (MSC-EVs) offer a promising cell-free alternative to MSC therapy, containing various biomolecules like proteins, mRNAs, and miRNAs that can be transferred to target cells [1–3]. MSC-EVs demonstrate therapeutic potential in animal disease models, mirroring the effects of their stem cell counterparts while providing advantages such as lower immunogenicity and the ability to cross biological barriers [4]. They are crucial in tissue repair, immunosuppression, and anti-inflammatory processes through paracrine actions [5].

Recent research indicates that dental stem cells (DSC) possess greater plasticity, proliferation, and immunomodulatory capacity than bone marrow-derived stem cells (BMSC) [6]. Dental pulp stem cells (DPSC) and periodontal ligament stem cells (PDLSC) can differentiate into bone, adipose tissue, cartilage, muscle, liver, and neural cells [7, 8]. Dental pulp stem cell-derived EVs (DP-EVs) promote bone regeneration in rat calvarial defects and enhance osteogenic differentiation of adipose-derived stem cells via the MAPK signaling pathway [9, 10]. They also exhibit proangiogenic effects, stimulating endothelial cell proliferation, migration, and formation, and support rapid neovascularization for dental pulp regeneration when combined with fibrin gel [11, 12]. DP-EVs carry bioactive molecules that contribute to their anti-inflammatory, osteo/odontogenic, angiogenic, and immunomodulatory functions [13, 14]. Similarly, periodontal ligament stem cell-derived EVs (PD-EVs) share these functional attributes [13]. When combined with collagen membranes, they enhance bone regeneration and regulate osteogenic and odontogenic differentiation via the miR-758-5p/LMBR1/BMP2/4 axis [15, 16]. PD-EVs also promote stress-induced osteogenesis, potentially increasing orthodontic tooth movement efficiency [17]. Despite the promising applications of MSC- and DSC-EVs, challenges remain in producing consistent EV preparations. MSC cultures are often heterogeneous, with senescence and differentiation affecting their phenotype and, consequently, the quality of the derived EVs. Since EVs reflect the molecular profile of their parent cells, maintaining uniformity in cell culture conditions is critical to ensuring the functional reliability of EVs. This variability underscores the need for standardized protocols and systematic studies to evaluate the therapeutic potential of EVs from different cell sources [18].

The molecular characteristics of EVs derived from various stem cell types are not well-defined, and existing research is often limited by variations in donor sources and culture conditions [19, 20]. Such inconsistencies hinder the identification of the most suitable cell source for specific therapeutic applications. Addressing this gap, the present study systematically compares EVs derived from DPSC and PDLSC, isolated from the same donors and cultured under identical conditions. By employing a standardized isolation protocol and including gingival fibroblast-derived EVs (GF-EVs) as a comparator, we aim to provide the first detailed characterization of EVs from different stem cell types sourced from the same individual. This work highlights potential functional differences and advances our understanding of the therapeutic applicability of stem cell-derived EVs.

## Materials and methods

### Cell culture

Healthy human third molars and premolars, extracted during routine orthodontic or prophylactic procedures, were collected by the Department of Oral and Maxillofacial Surgery (RWTH Aachen University Hospital, Aachen, Germany), as approved by the local ethics committee (EK 458/21). Periodontal ligament tissue was isolated from the root’s middle third, and dental pulp tissue was dissected from the pulp cavity and root canal, then minced into 1-2 mm segments. Both tissues were digested separately with collagenase I and dispase II (Gibco, Carlsbad, CA, USA). PDLSC and DPSC were cultured in Alpha Modified Eagle’s medium (α-MEM) with 15% FBS and 1% antibiotics/antimycotics (Gibco, Carlsbad, CA, USA) and identified by flow cytometry (BD biosciences, San Jose, CA, USA) using markers such as CD11b, CD34, CD45, CD73, CD79, CD90, CD105 and HLA-DR. Cells up to passage 6 were used. Immortalized human GF were obtained commercially from Innoprot (Derio, Bizkaia Spain). The cells have been developed by immortalizing primary human gingival fibroblasts with SV40 Large T antigen. Human umbilical vein endothelial cells (HUVECs) were sourced from RWTH Aachen University’s Centralized Biomaterial Bank (cBMB) according to its regulations, following the RWTH Aachen University, Medical Faculty Ethics Committee’s approval (cBMB project number 323). The lumen of the umbilical cord vein was soaked with collagenase (Thermofischer Scientific, Waltham, MA, USA) for 30 minutes at 37 °C. After washing with PBS and collagenase removal, the cells were cultured in cell culture flasks and supplied with an endothelial basal medium (EGM-2, Promocell, Heidelberg, Germany).

### EVs isolation

DPSC, PDLSC and GF were cultured in α-MEM until reaching 70% confluence. The medium was then replaced with exosome-depleted medium, and cells were incubated until 95% confluence. EVs were isolated from the culture supernatant using differential centrifugation, following established protocols [21, 22]. The process involved an initial centrifugation at 300 × g for 10 minutes to remove cells, followed by centrifugation at 2 000 × g for 20 minutes. The supernatant was then centrifuged at 10 000 × g for 40 minutes, filtered through 0.2 um diameter filter, and subjected to ultracentrifugation at 110 000 × g for 70 minutes. The resulting pellet was washed with PBS and centrifuged again at 110 000 × g. The temperature was maintained at 4°C throughout all centrifugation processes. Ultracentrifugation was performed using a Beckman L-90K ultracentrifuge and a matching SW 32 Ti rotor. The EV pellet was finally resuspended in 100 µl PBS and stored at −80°C. CM without EVs was also stored and served as a control.

### Characterization of EVs

The presence of EVs was confirmed by transmission electron microscope (TEM). The pelleted EVs were suspended in double-distilled water and applied dropwise onto formvar carbon-coated nickel grids. The grids were air-dried, stained with 2% uranyl acetate and viewed under a Zeiss LEO 906E transmission electron microscope (Zeiss, Jena, Germany).

The size distribution and concentration of EVs were determined by NTA, using NanoSight NS300 (Malvern Instruments, Worcestershire, UK) equipped with a blue laser (488nm, 70mW). EV samples were diluted 1:100 in filtered PBS prior to NTA. Data was performed with NTA 3.0 software (Malvern Instruments, Worcestershire, UK).

The total protein content of EVs were detected using Rapid Gold BCA Protein Assay Kit (Thermo Fisher Scientific, Waltham, MA, USA) according to manufacturer instructions.

EV surface markers were detected by Imaging Flow Cytometry (IFCM) described in Ricklefs et al [23]. In short, EVs were stained in filtered PBS containing 8% EV-free FCS supplemented with protease-inhibitor and phosphatase-inhibitor. Antibodies used to stain EVs were PE-conjugated anti-CD9 (Biolegend, clone HI9a, 20 μg/mL, diluted 1:30), PacBlue-conjugated anti-CD63 (Biolegend, clone H5C6, 40 μg/ml), FITC-conjugated anti-CD81 (Biolegend, clone 5A6, 40 μg/ml) and an isotype control (MOPC-21, 500 μg/ml). Staining was performed by incubating EVs with antibodies for 45 minutes at room temperature in the dark. Samples were then washed with 2% EV-free FCS, resuspended, and analyzed using the AMNIS ImageStream^x^ Mark II Flow Cytometer (AMNIS/Millipore, Seattle). Data analysis was conducted using IDEAS software version 6.2.

For the phenotypical characterization, the MACSPlex Exosome kit (Miltenyi Biotec, Bergisch Gladbach, Germany) was used. This bead-based assay allows the simultaneous detection of 37 surface markers. In brief, EVs were incubated overnight with antibody-coated capture beads, washed, and incubated with antibodies provided in the kit. Analyzation of the bead-bound EVs was done using FACSCanto II (BD Biosciences, La Jolla, USA). Quantifying and comparing the expression of different EV markers across samples allowed the assessment of variations in EV populations.

### EVs internalization assay

EVs were labelled with PKH26 fluorescent dye. Excess dye was removed through washing and centrifugation. HUVECs were cultured until 50% confluence, then media was changed and supplemented with labelled EVs at 1.2 × 10^10^/ml. Cells were fixed at 1, 3, 6, 12, and 24 hours, then stained with phalloidin 488 and DAPI. EV uptake was visualized and quantified using Nikon inverted microscope (Nikon Eclipse Ti Series, Japan) and ImageJ software (Adobe Systems, San Jose, CA, USA), measuring fluorescence intensity per cell to determine the internalization of EVs.

### Cell proliferation assay

HUVECs were seeded into a 96-well plate with a density of 5 000 cells per well to ensures uniform distribution and cultured overnight in EGM-2 medium. The following day, the medium was replaced with fresh medium containing EVs at a concentration of 1.2 × 10^10^/ml. EV-treated HUVECs were incubated at 37°C for various time points (6, 12, 24, 48, and 72 hours). To evaluate cell viability, the XTT (Sigma Aldrich, USA) assay was performed. XTT reagent was prepared according to the manufacturer’s instructions. After the specified incubation times, 50 µL of the prepared XTT mixture was added to each well. The absorbance was measured at 450 nm using a microplate reader. The results from EV-treated cells were compared to untreated control cells to assess the impact of EVs on cell viability.

### Endothelial cell tube formation assay

To evaluate the effect of EVs on vascularization, five distinct HUVECs/GF 3D-co-culture systems were established using a fibrin gel matrix [24]. These systems included gels containing DP-EVs, PD-EVs, GF-EVs, CM without EVs, and pure gels as a negative control. The final concentration of EVs in the gels was standardized to 1.2 × 10^10^ particles/mL. Fibrin gels were prepared by mixing fibrinogen (Human Fibrinogen, Enzyme Research Laboratories, US) with HUVECs and GF in a 1:1 ratio, achieving a final cell density of 14.5 million cells/ml. Polymerization was initiated by adding thrombin and CaCl₂ within a 3D cell culture chip (AIM BIOTECH, Singapore). The cultures were incubated, with medium (EGM-2) replaced daily to maintain optimal conditions. After seven days, samples were fixed and stained for CD31 (PECAM-1, Sigma) to visualize vascular structures. Imaging was performed using a confocal microscope (Zeiss LSM 980 Airyscan 2). Quantitative analysis of vascular volume, length, and diameter was conducted using Imaris software (version 10.20).

### miRNA expression assay

RNA was extracted from 60 µL of EVs samples using the Maxwell RSC miRNA Plasma and Serum Kit (Promega, Madison, WI, USA), following the manufacturer’s protocol. The RNA concentration was measured with a Promega Quantus Fluorometer (Promega, Madison, WI, USA). Sequencing libraries were prepared using the QIAseq miRNA UDI Library Kit (QIAGEN, Hilden, Germany), following the manufacturer’s protocol. To monitor the library preparation process and ensure the accuracy of miRNA detection, QIAseq miRNA Library QC Spike-ins were added to each sample. The size distribution of the libraries was assessed using the Agilent TapeStation (Agilent Technologies, Santa Clara, CA, USA). The average library size was confirmed to fall within the expected range for miRNA. Library concentrations were then quantified using the Promega Quantus Fluorometer to ensure appropriate loading for sequencing.

Sequencing was performed on an Illumina NextSeq 500/550 system using the High Output Kit v2.5 (75 cycles) (Illumina, San Diego, CA, USA). Libraries were sequenced in single-end mode for 72 cycles, with a 1% PhiX spike-in used as an internal control to monitor sequencing quality and performance. FASTQ files were generated with bcl2fastq (Illumina). To facilitate reproducible analysis, samples were processed using the publicly available nf-core/smRNAseq pipeline version 1.1.0 implemented in Nextflow 21.10.6 using Docker 20.10.12 with the minimal command [25–27]. All analysis was performed using custom scripts in R version 4.1.1 using the DESeq2 v.1.32.0 framework [28].

### EVs proteomics analysis

To isolate proteins from EV the SP3 protocol was used [29], and disulfide bonds were first reduced with dithiothreitol, followed by alkylation with iodoacetamide in the dark. The samples were then dissolved in 70% acetonitrile and incubated with carboxylate-modified magnetic beads for protein binding. After washing with acetonitrile and ethanol, trypsin digestion occured overnight. The digested peptides were then eluted and stored for further use. For chromatographic separation, a UHPLC system was used with a two-buffer system (buffer A with 0.1% formic acid in water and buffer B with 0.1% formic acid in acetonitrile). The peptides were separated using a C18 reversed-phase column with an 80-minute gradient. In the mass spectrometry analysis, the Exploris 480 instrument performs data-dependent acquisition with nano-ESI ionization. The highest-intensity peptides are fragmented, and MS2 scanning was carried out. Data analysis was performed with Proteome Discoverer software using the Sequest algorithm, and proteins were identified with strict false discovery rate (FDR) criteria. Quantification was achieved using the Minora algorithm, and protein abundances were log2-transformed and normalized.

### Statistical analysis

Statistical analyses were performed using GraphPad Prism (GraphPad Software, San Diego, USA). The normality of the data was assessed using the Shapiro-Wilk test. F-test was performed for the comparison of variances. Statistical comparisons were performed using one-way ANOVA with Tukey’s post-hoc test for the normally distributed data. Nonparametric analysis of non-normally distributed data was conducted using Kruskal-Wallis test. A *p*-value of ≤ 0.05 was considered statistically significant.

## Results

### Isolated DPCS and PDLSC express surface marker characteristics of mesenchymal stem cells

Flow cytometry analysis showed that the majority of the isolated DPSC and PDLSC expressed key mesenchymal stem cell markers, including CD73, CD105, and CD90, with over 94% of the cells testing positive for these markers. In contrast, markers associated with the hematopoietic lineage (CD45, CD34, CD11b, CD79a, HLA-DR) were expressed in less than 4% of the population. These results confirm that the isolated DPSC and PDLSC exhibit typical characteristics of mesenchymal stem cells (Tab. 1). The immortalised GF expressed the investigated surface markers to the same extent.

**Tab. 1.**
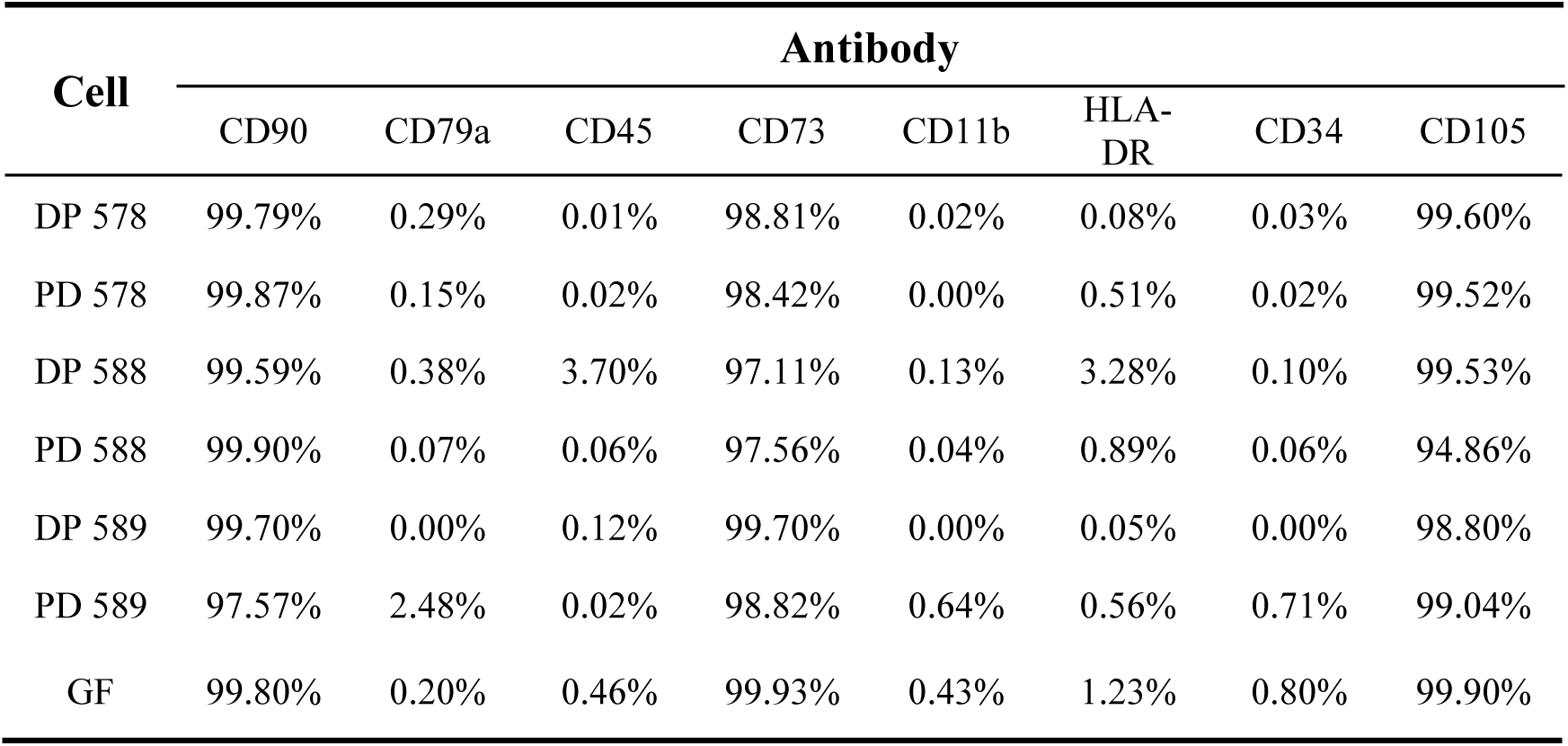
Cell surface marker expression for stem cell characterization. DP indicates dental pulp stem cells, PD periodontal ligament stem cells, the numbers represent the donor.

| Cell | Antibody |  |  |  |  |  |  |  |
| --- | --- | --- | --- | --- | --- | --- | --- | --- |
|  | CD90 | CD79a | CD45 | CD73 | CD11b | HLA-DR | CD34 | CD105 |
| DP 578 | 99.79% | 0.29% | 0.01% | 98.81% | 0.02% | 0.08% | 0.03% | 99.60% |
| PD 578 | 99.87% | 0.15% | 0.02% | 98.42% | 0.00% | 0.51% | 0.02% | 99.52% |
| DP 588 | 99.59% | 0.38% | 3.70% | 97.11% | 0.13% | 3.28% | 0.10% | 99.53% |
| PD 588 | 99.90% | 0.07% | 0.06% | 97.56% | 0.04% | 0.89% | 0.06% | 94.86% |
| DP 589 | 99.70% | 0.00% | 0.12% | 99.70% | 0.00% | 0.05% | 0.00% | 98.80% |
| PD 589 | 97.57% | 2.48% | 0.02% | 98.82% | 0.64% | 0.56% | 0.71% | 99.04% |
| GF | 99.80% | 0.20% | 0.46% | 99.93% | 0.43% | 1.23% | 0.80% | 99.90% |

After 79 times of EV isolation from DPSC and PDLSC lines of a total of 10 donors, and immortalized GF cell line in different passages, we could not find any correlation neither for the particle number depending on the cell number nor for the whole protein content depending on the EV particle number (Fig. 1A/B). For the reasons mentioned above and the fact that contamination with non-vesicular proteins cannot be excluded when FBS is used in cell culture, we decided to use the EVs particle number as the dosage parameter in the following experiments. After generating scatter plots, performing linear regression analysis, and setting a 95% confidence interval (Figure 1A/B). R^2^ is < 0.1, suggesting a weak or negligible correlation between the variables.

**Fig. 1.**
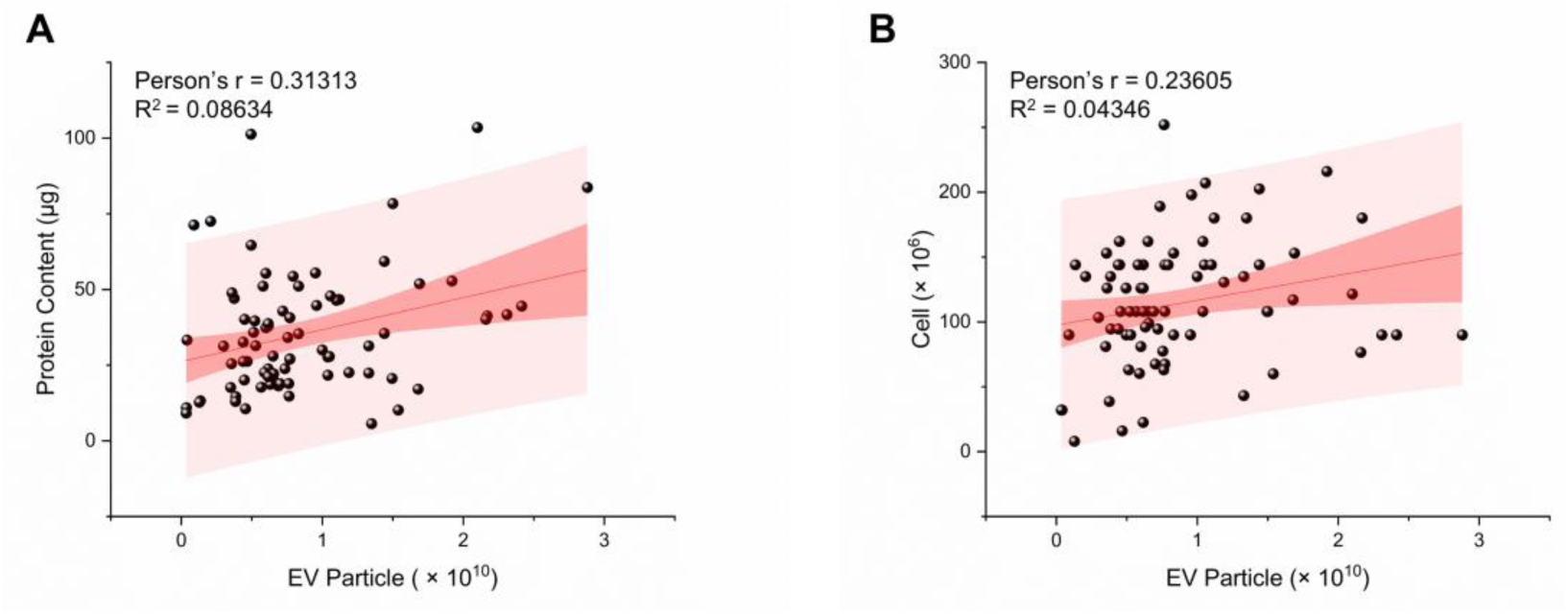
Correlation between cell number, EV particle number and protein content. (**A**) Correlation between the number of EV particles and protein content. (B) Correlation of the number of EV particles with the number of cells of origin. R^2^ < 0.1, n = 79.

### Characteristics of EVs

TEM revealed a characteristic cup-shaped morphology of the EVs. The size distribution ranged from 50 to 200 nm, with a peak size of around 120-140 nm for all three different EVs (Fig. 2A). IFCM confirmed the presence of key surface markers CD9, CD63, and CD81 (Fig. 2B).

**Fig. 2.**
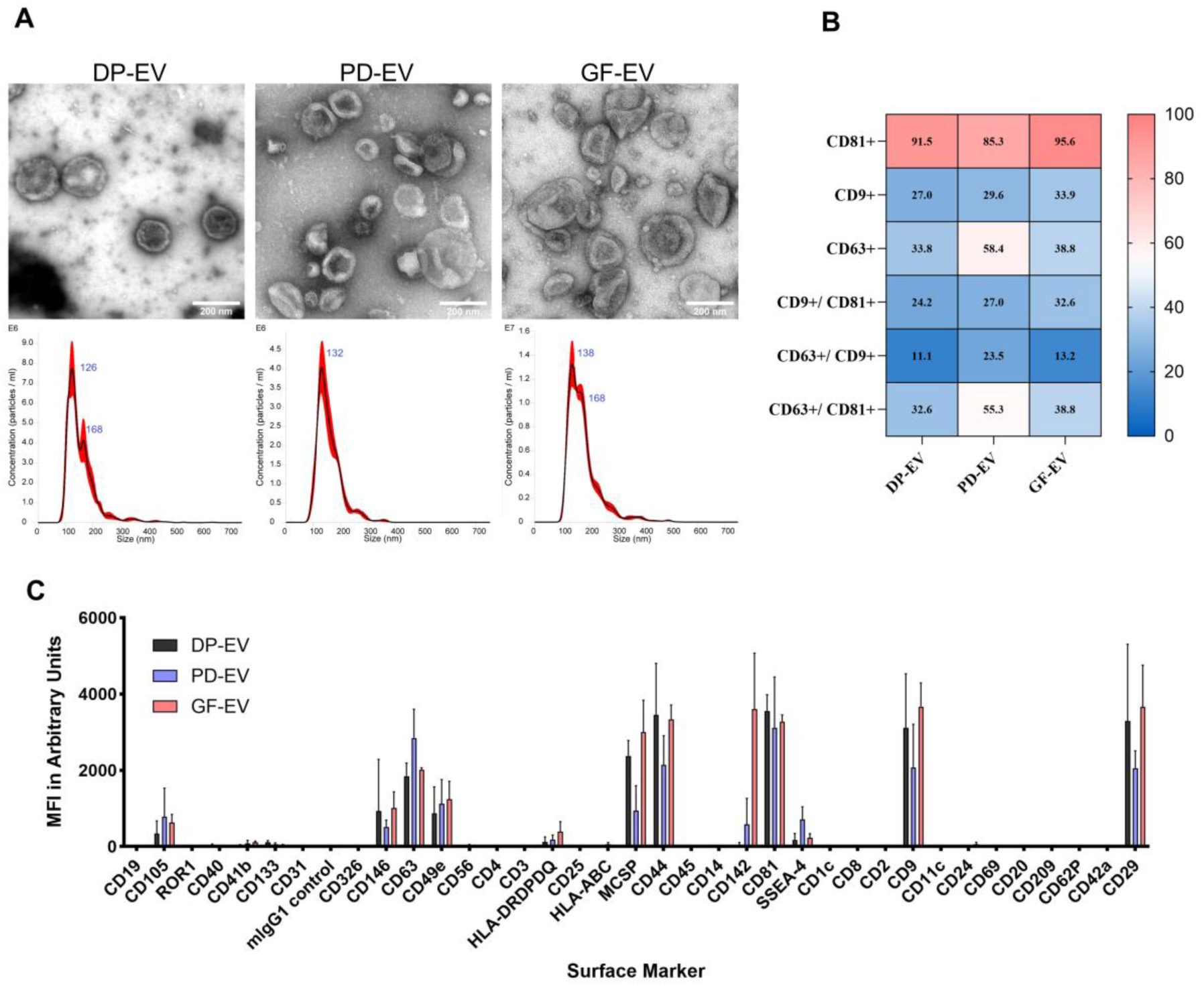
Characterization of EVs by TEM, NTA, and surface marker expression. (**A**) TEM revealed the characteristic cup-shaped morphology of EVs. NTA confirmed a similar size distribution across all three EV preparations, with an average size of approximately 130 nm. (**B**) Imaging flow cytometry showed that all EV samples expressed tetraspanins, with CD81 exhibiting particularly high expression. EVs derived from PDLSC displayed a notably strong combined expression of CD63 and CD81. (**C**) Bead-based multiplex assay further highlighted the expression of common EV surface markers, underscoring the consistent marker profile across the samples.

Multiplex bead-based assay analysis shows high levels of typical EV surface markers (CD9, CD81, and CD63) as well as other relevant antigens (e.g., CD105, CD44, CD29) were detected in the three different EV samples. Other markers abundantly detected comprised mainly CD146, CD49e, and MCSP. Markers detected at intermediate to low-positive fluorescence intensity were HLA-DR/DP/DQ and SSEA-4. The platelet tissue factor CD142 was abundantly detected in GF-EVs, slightly detected in PD-EVs, and, in contrast, exhibited a quite low detection value in DP-EVs (Fig. 2C).

### Internalization of EVs

EV uptake assay demonstrated that PKH-26 labelled EVs were effectively internalized by HUVECs in a time-dependent manner. As shown by the appearance of red dots (Fig. 3A), fluorescence microscopy confirmed the HUVECs internalized a small amount of EVs at 1 h, the red fluorescence intensity inside cells increased significantly at 3, 6 and 12 hours. Starting from 12 hours, the internalization of all three EV groups was significant compared with the control group. The internalization of PD-EVs declined at 24 hours, while DP-EVs and GF-EVs continued to increase until 24 hours. As endocytosis increased, EVs were observed aggregated toward the nucleus of HUVECs (Fig. 3A). From 6 hours onwards, the internalization of GF-EVs significantly higher than PD-EVs. (Fig. 3B). The time-dependent internalization of DP-EVs was statistically significant higher compared to the other EV types at 6 hours and 12 hours.

**Fig. 3.**
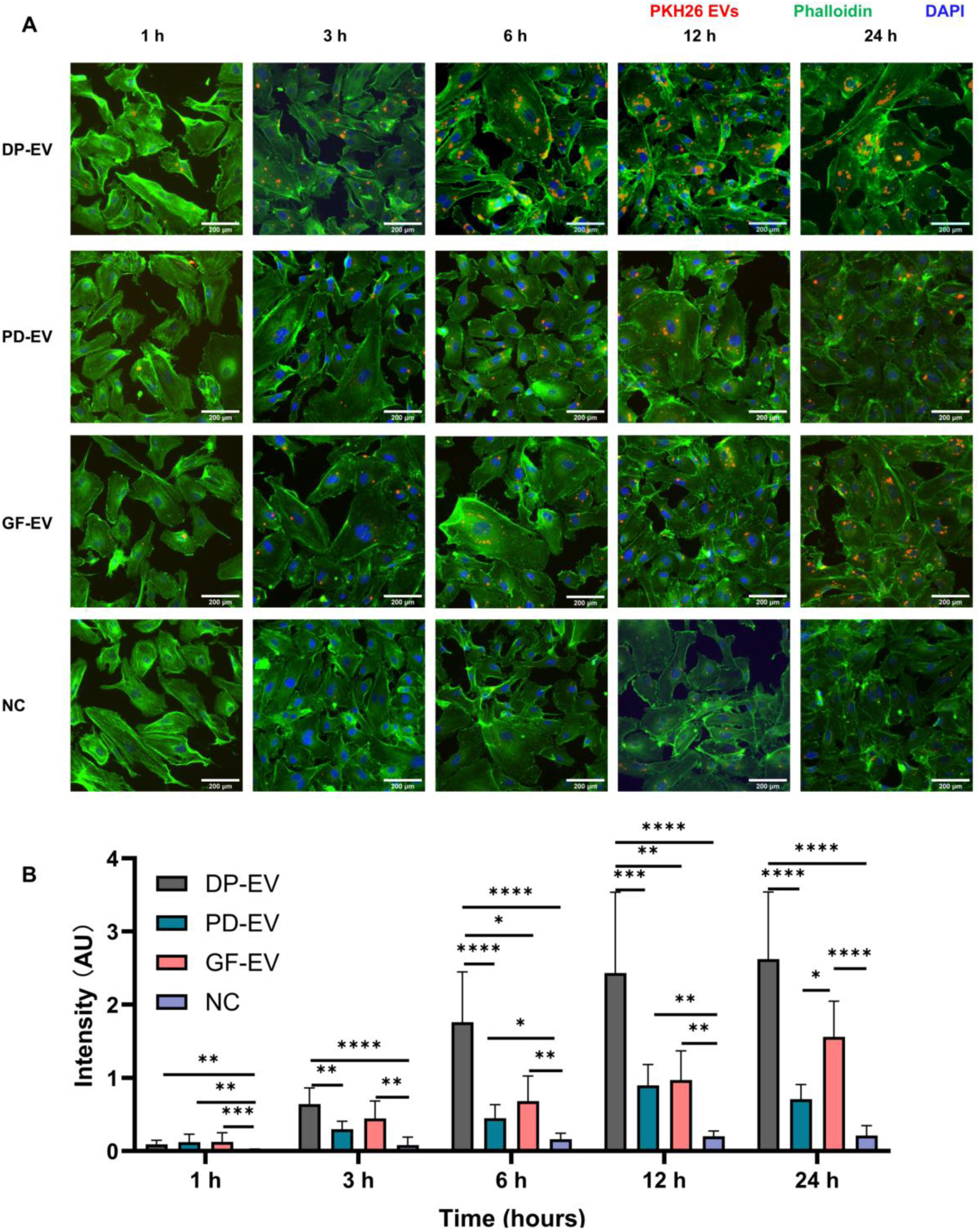
Uptake of EVs by HUVECs. (**A**) Time dependent of different EV internalization. HUVECs were incubated with of PKH26 (red) labelled PD-EVs, DP-EVs, GF-EVs and PBS as EV free control for 1, 3, 6, 12 and 24 hours. HUVECs were stained for actin (phalloidin, green) and nuclei (DAPI, blue). Starting from 3 h after incubation, the majority of internalized EVs were detected in the perinuclear region of cells. Scale bar, 200 μm. (**B**) Internalized EVs were quantified by means of fluorescence intensity. Kruskal-Wallis test, \**p* < 0.05, \*\**p* < 0.01, \*\*\*\**p* < 0.0001, n = 3.

### Endothelial cell viability assay

The XTT cell viability assay results revealed the effects of the three different EVs and conditioned medium on HUVECs viability over time (Fig. 4). Unlike the EV internalization experiment, where cells gradually accumulated EVs and significant differences between groups were observed within the first 24 hours, no significant differences in cell viability were detected between the experimental and control groups during this initial period. Statistical differences began to emerge at 48 and 72 hours. A fluctuation in cell viability was observed in all groups during the first 24 hours. From 24 to 72 hours, cell viability in the EV-treated groups gradually increased, whereas viability in the control group steadily declined.

**Fig. 4.**
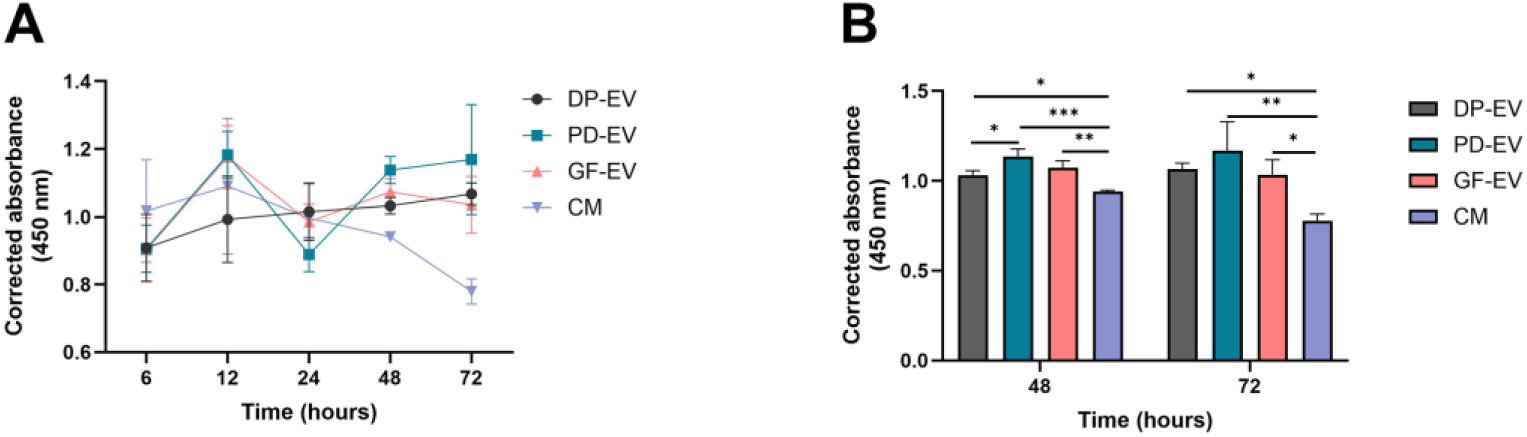
Results of XTT viability assay. (**A**) Effects of DP-EV, PD-EV, GF-EV, and CM (supernatant after ultracentrifugation step of EV isolation) on HUVECs viability over 72 hours. (**B**) Statistical evaluation of cell viability at 48 and 72 hours. Data were analyzed using one-way ANOVA with Tukey’s post-hoc test, \**p* < 0.05, \*\**p* < 0.01, n = 3.

### Tube formation in fibrin gels

To investigate the proangiogenic effect of EVs, we cultured HUVECs together with GF as supporting cells in a 3D environment in fibrin gel. Confocal microscopy showed well-grown neovascular structures on culture day 7 (Fig. 5A). Quantification of 3D tube formation showed that DP-EVs and GF-EVs significantly improved vascularization compared to the control group, as evidenced by increased vessel length and volume. In contrast, the CM and PD-EVs group exhibited lower neovascularization activity, indicating the superior potential of DPSC- and GF-derived EVs in promoting vessel formation. No significant differences were found in the diameter of the formed tubular structures (Fig. 5B).

**Fig. 5.**
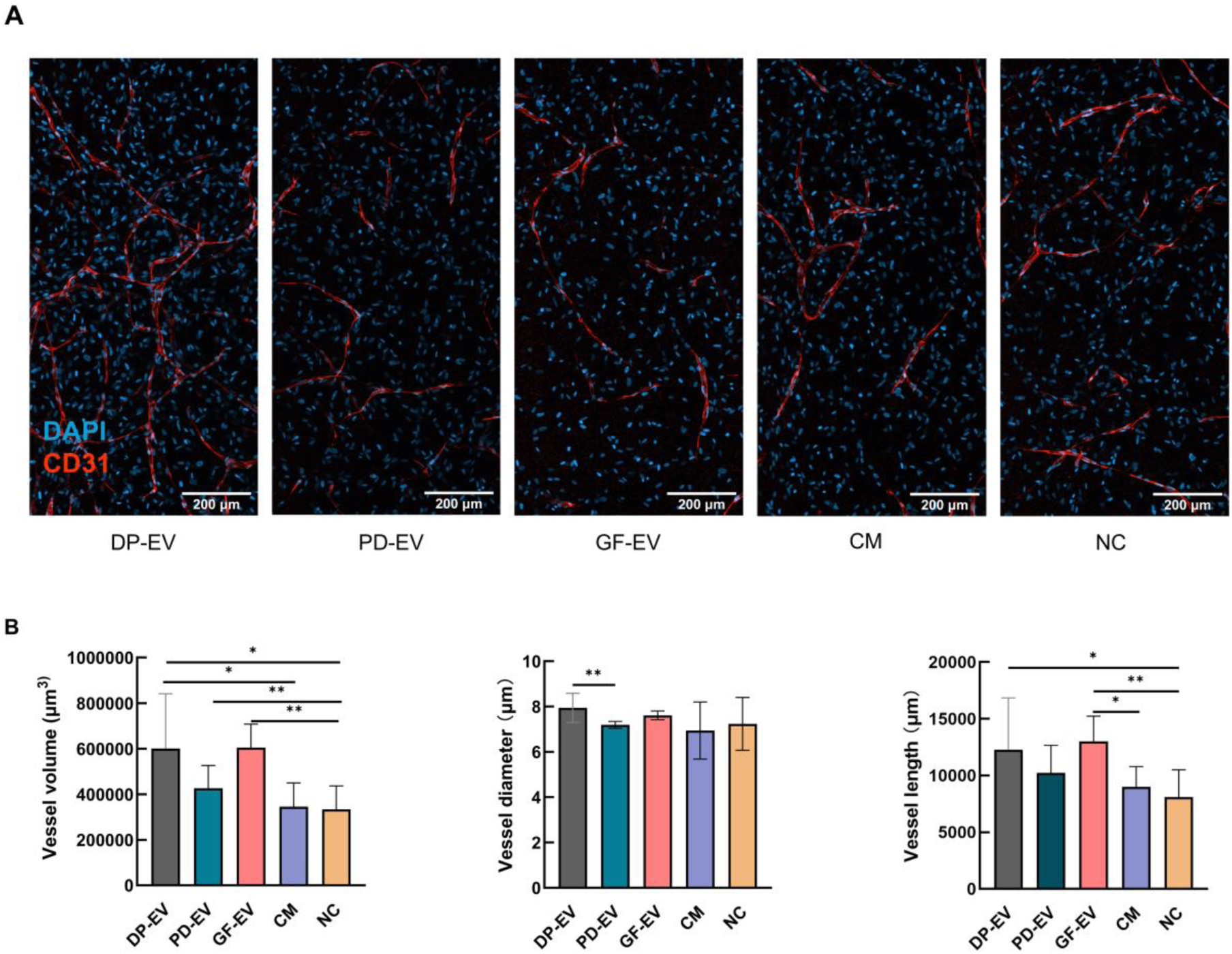
Tube formation assay of HUVECs-GF co-culture within 3D cell culture chip. (**A**) Laser scanning confocal microscopy image of CD31 staining after 7 days of cultivation treat with DP-EV, PD-EV, GF-EV, CM and PBS as control. Scale bar: 200 μm. (**B**) Statistical evaluation of volume (Kruskal-Wallis test), diameter (Kruskal-Wallis test), and length of vascular structure (One-way ANOVA with Tukey’s post-hoc test). \**p* < 0.05, \*\**p* < 0.01; n = 3.

### EV-miRNA Sequencing

To characterize the miRNA profile of the herein investigated EVs, we conducted a miRNA sequencing analysis (Fig. 6). The Venn diagram revealed that 86 of the 100 most frequently expressed miRNAs were shared across all three EV groups (Fig. 6A). Eleven miRNAs were exclusively present in GF-EVs, four in PD-EVs, and only one specific for DP-EVs. Notably, the heatmap of the top 100 miRNAs highlighted distinct expression patterns (Fig. 6B). However, the 20 most abundant miRNAs showed a strong overlap across the groups (Fig. 6C). Further principal component analysis (PCA) showed a clear clustering of the analyzed samples (Fig. 6D). Remarkably, the clustering was donor-dependent, whereas the origin cell type of the EVs did not play a primary role. Interestingly, the miRNA data for GF-EVs did not cluster, despite being derived from the same cell line across only two different passages. To explore the functional implications of the miRNA profiles, we analyzed the enrichment of KEGG signaling pathways for the 20 most abundantly expressed miRNAs (Fig. 6E). The results show that the signaling pathways are mainly involved in the process of transcriptional misregulation in cancer, progesterone oocyte maturation, microRNAs in cancer, regulation of stem cell pluripotency, P53, PI3K-Akt, FoxO and TGF-beta signaling pathways.

**Fig. 6.**
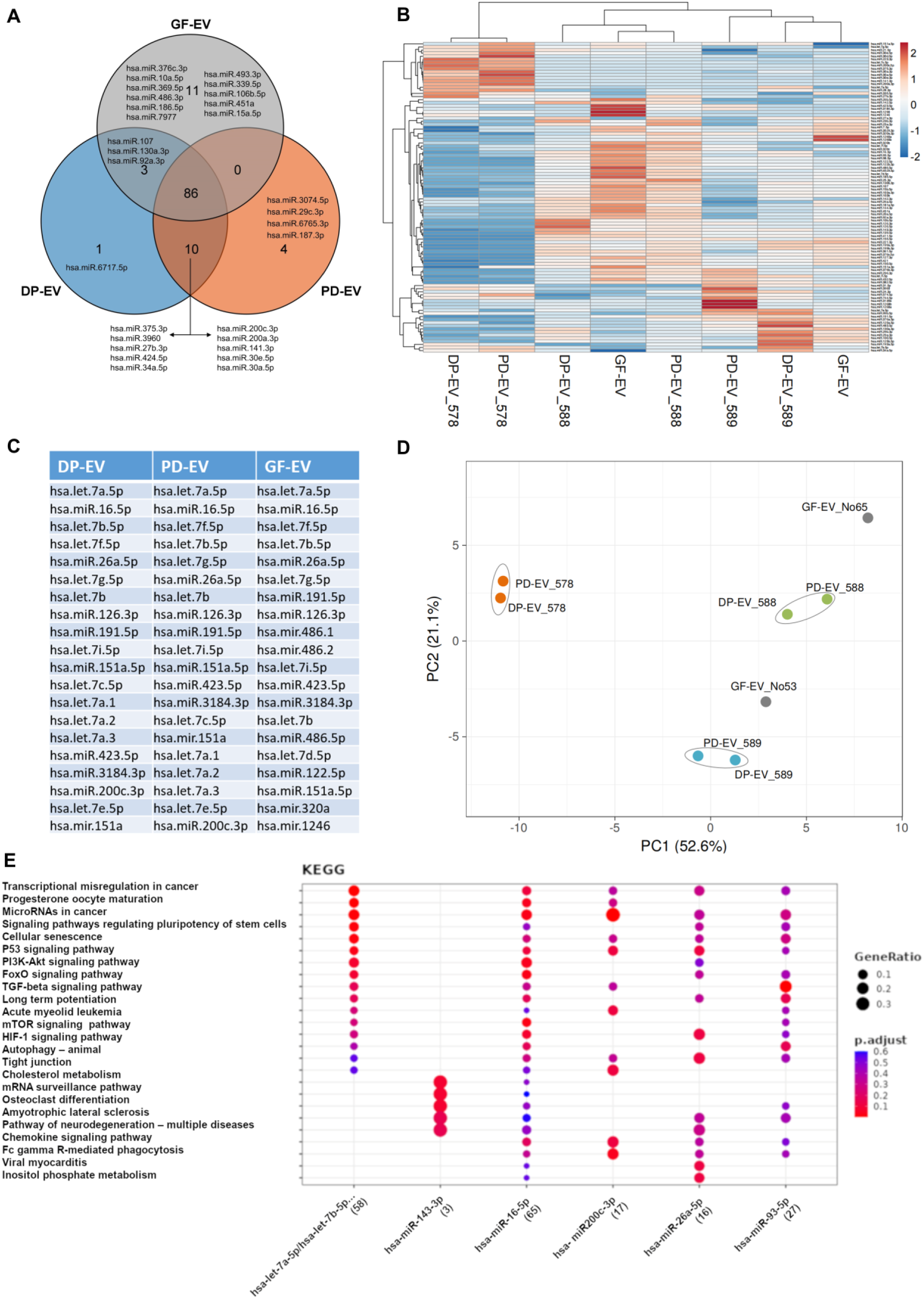
Differentially expressed miRNAs in DP-EV, PD-EV and GF-EV. (**A**) Venn diagram comparing the top 100 differentially expressed miRNAs in DP-EVs, PD-EVs, and GF-EVs. (**B**) Hierarchical clustering analysis: heat map showing the differentially expressed miRNAs by studied groups plotted using ClustVis (correlation distance measure and average linkage function). (**C**) Table of the top 20 miRNAs expressed in DP-EV, PD-EV and GF-EV. (**D**) PCA of the miRNA profiling of three samples. The X and Y axes show principal component 1 and principal component 2, which explain 52.6% and 21.1% of the total variance, respectively. (**E**) Enrichment analysis of top 20 miRNAs performed MIENTURNET (MicroRNA ENrichment TURned NETwork) web tool selecting the KEGG pathway [30].

### Proteomics

As shown in Figure 7A, among the 20 most frequently expressed proteins, 12 proteins were present in all 3 EVs. These included lactadherin (MFGE8), filamin-A (FLNA), Ras-related protein Rap-1b (RAP1B), and moesin (MSN). Among the top 20, 5 proteins were exclusive to the DP-EVs. These included for example CD44 and eukaryotic translation elongation factor 1 alpha 1 (EEF1A1). Three proteins were exclusive to the PD-EVs. These included tubulin alpha 4a (TUBA4A). Only two proteins were exclusive to GF-EVs, alpha-2-macroglobulin (A2M) and integrin beta-1 (ITGB1). Among the top 100 occurring proteins, the agreement was as high as 99%. Despite the large overlap, the heatmap shows distinctive patterns in terms of abundance (Fig. 7B). To observe the functional enrichment of the top 100 EV proteins, g: Profiler was used for the analysis [31]. The parameters for the enrichment analysis were as follows: H. sapiens was chosen as organism. The gene ontology analyses (GO Molecular Function (GO: MF), GO Biological Process (GO: BP) and GO Cellular Component (GO: CC)) were performed sequentially. The KEGG, Reactome and WikiPathways databases were used for biological pathways. GO enrichment analysis revealed that 65, 318 and 115 GO terms were enriched in MF, BP and CC, respectively. Classification analysis revealed that GO: MF mainly focused on cell adhesion molecule binding, GO: BP on wound healing, and GO: CC on cell-substrate junction and EVs. A total of 28, 76 and 17 pathways were enriched in KEGG, REAC and WP, which were mainly related to focal adhesion, response to increased platelet cytosolic Ca2+ and focal pathways, respectively (Fig. 7C). The table lists 10 selected terms that were associated with high statistical significance (Fig. 7D).

**Fig. 7.**
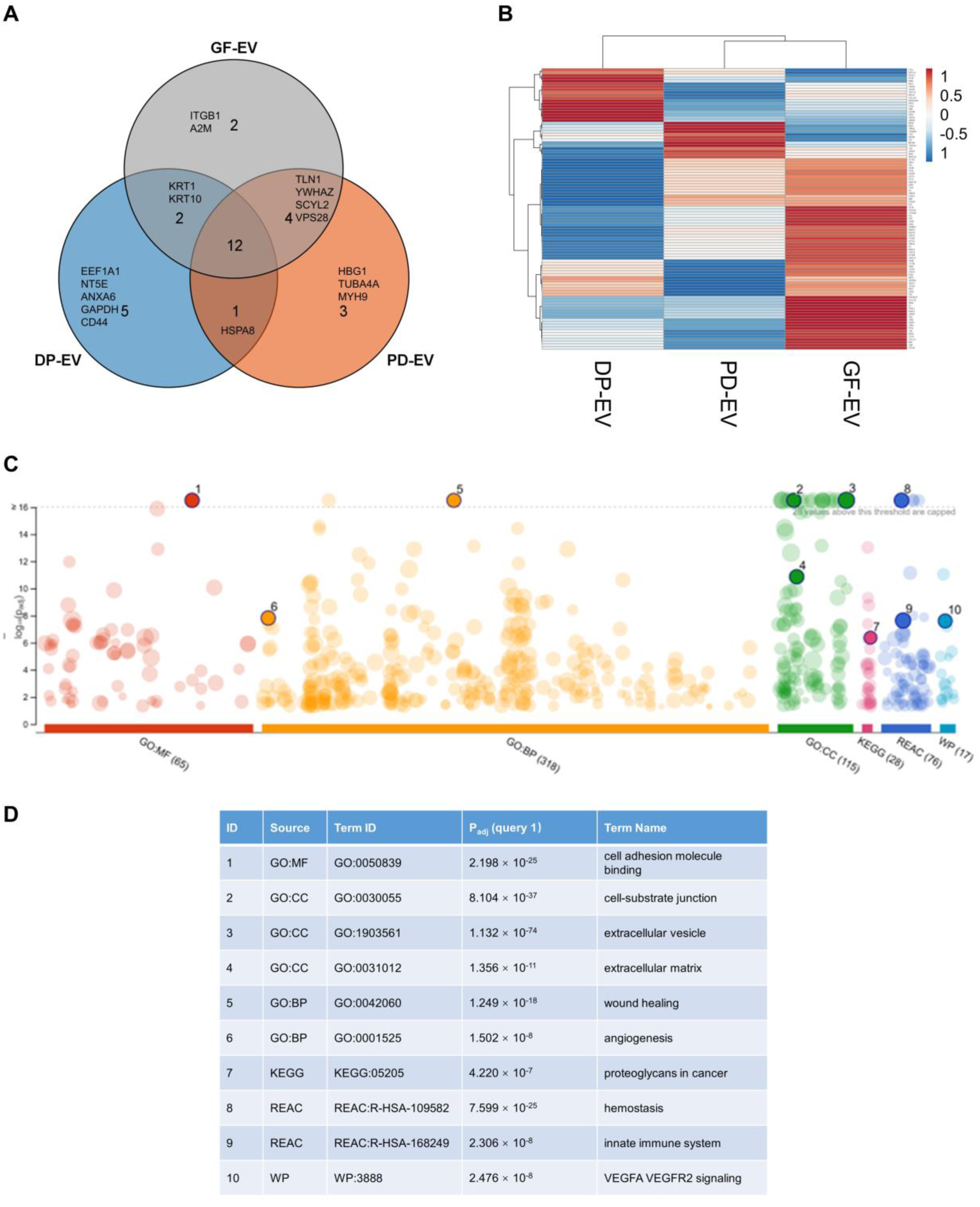
Differentially expressed proteins in DP-EV, PD-EV and GF-EV. (**A**) Venn diagram comparing the top 20 expressed proteins in DP-EVs, PD-EVs, and GF-EVs. (**B**) Differentially expressed proteins were visualized using a heat map. (**C**) Bioinformatics analysis of the top 100 EV proteins performed by g:Profiler. A specific organism was chosen H. sapiens (human). GO analyses of molecular function (MF), biological process (BP), and cellular component (CC) were carried out sequentially. The biological pathways used were the KEGG, Reactome, and WikiPathways databases. (**D**) Table of part pathways in g:Profiler enrichment analysis.

## Discussion

The present study explored EVs derived from various odontogenic cell lines to assess their potential for regenerative medicine. Teeth are an accessible and non-invasive source of several different stem cell types and therefore provide an excellent opportunity to compare tissue-specific and donor-dependent variations. To our knowledge, the present work represents the first analysis of EVs from stem cells of different tissues within a single individual. Previous studies have shown that DPSC and PDLSC, both originating from the neural crest, exhibit distinct biological properties influenced by their niches. DPSC exhibits higher osteogenic potential and a stronger response to osteogenic stimuli, while PDLSC excels in mineralization and migration [32, 33]. Both cell types express mesenchymal stem cell markers, but DPSC show higher expression of stemness-related markers, suggesting that they may be more primitive [34]. PDLSC have a higher neuronal differentiation rate than DPSC and show better therapeutic effects in a rat model [35]. DPSC exhibit stronger immunomodulatory capabilities, expressing immunostimulatory ligands when primed with IFNγ [36]. The proliferation rate of PDLSC is generally lower than that of DPCS [37]. These differences underscore their unique roles and applications in regenerative medicine. GF are critical to maintaining oral health, contributing to tissue structure, immune regulation, and extracellular matrix production. GF act as sentinel cells, responding to pathogens by producing inflammatory mediators [38]. Their unique properties, originating from the neural crest and strong immunomodulatory functions, make GF extremely attractive for therapeutic applications [39, 40]. GF contributes to wound healing and tissue repair but can also promote chronic inflammation in periodontitis [41]. In contrast to stem cells, GF are stable, highly proliferative, and long-lived, making them suitable for research and testing in regenerative applications.

Based on the reasons described above, we have selected these cell lines. The cells cultivated showed the usual expression patterns of the surface antigens of adult mesenchymal stem cells. Interestingly, the GF showed a similar expression pattern. However, GF are heterogeneous, with subpopulations exhibiting various surface markers, including CD59, CD99, CD9, CD95, CD55, CD63, CD26, CD117, CD71, and CD86 and functions [42, 43]. Furthermore, flow cytometric analysis found that CD73 and CD90 are highly expressed by human PDLSC and GF [44].

EVs were isolated from the supernatant of the three cell lines by differential ultracentrifugation, which is a widely used method for isolating EVs from various biological fluids. This technique involves sequential centrifugation steps to separate EVs based on size and density. However, the method has limitations, including co-isolation of contaminants and variability in yield between laboratories [45]. Factors such as rotor type, centrifugation time, and g-force significantly influence EV isolation efficiency [46]. Despite these challenges, ultracentrifugation remains a valuable tool for EV isolation, enabling subsequent characterization through techniques such as nanoparticle tracking analysis (NTA), transmission electron microscopy, western blotting, and flow cytometry [47].

Consistent with the recommendation of the International Society for Extracellular Vesicles (ISEV), TEM and NTA showed that the EVs were small, approximately 120–140 nm, and expressed the standard markers CD9, CD63, and CD81. These tetraspanins, in addition to being widely used EV markers, are involved in EV biogenesis, cargo selection, and cellular targeting, which underlies their effect on distinct cellular processes under both physiological and pathological conditions [48]. In this context it is noteworthy that all three EV types showed a high expression of CD81+, a protein commonly abundant in brain cells and EVs [49]. CD63, known for its role in regulating EV secretion and its association with MSC [50], was significantly elevated in PD-EVs. The proportion of double positive CD81+/CD9+ was almost the same in all three EV types. However, the frequency of tetraspanins was comparable in DP-EVs and GF-EVs, while the PD-EVs differed.

To identify heterogeneity between the three EV types, we used a multiplex bead assay with a total of 37 surface markers. In addition to the marker proteins described above, CD105, CD146, CD49e, HLA-DR, CD44, SSEA-4 and CD29 were also detected. There were no major differences in the presence of these proteins on the different EVs. The only difference in expression was for CD142. This was highly expressed in GF-EVs, but only slightly in PD-EVs and almost not at all in PD-EVs. This is an interesting finding as CD142/tissue factor (TF)/coagulation factor 3 is a transmembrane glycoprotein that acts as a receptor for FVII/FVIIa and initiates the extrinsic coagulation pathway. The expression of CD142 is not cell-specific and TF is also found in fibroblasts [51].

Accurate quantification of EVs is crucial for studying their function. The correlation between particle counts and protein levels may be influenced by non-vesicular contaminants or protein aggregates, leading to inconsistencies in data. The ISEV guidelines for Minimal Information for Studies of Extracellular Vesicles (MISEV) does not recommend using protein concentration as surrogate for EV concentration [52]. However, a high particle-to-protein ratio generally indicates a purer EV preparation, while a lower ratio may imply the presence of non-vesicular protein contaminants. By analyzing 79 EV samples in our study, we found that, on average, 2.4 × 10¹⁰ EVs contained 100 μg of protein. However, no clear correlation was observed between EV particle count and protein content. EV samples from various cell sources can present different ratios of particle numbers to protein content, and certain purification methods may enrich for protein or particle contaminants, which skews this ratio. Standardization efforts like the MISEV guidelines have emphasized the need for rigorous isolation techniques to ensure that the EV protein content is accurately reflected in particle-based quantifications [52]. In this study, we strictly maintained consistent cell culture conditions and EV isolation method but still could not demonstrate a correlation between EV number and protein content. For this reason, we used particle number exclusively for standardisation in subsequent experiments. Interestingly, there was also no relationship between initial cell count and EV yield, despite controlled culture conditions and the use of low-passage cells. This variability highlights the difficulty of achieving consistent EV production, presenting a significant challenge for scaling up EV-based therapies and standardizing dosing in translational applications.

To evaluate the functional properties of EVs from three different cell sources, we first examined their uptake by endothelial cells. Using the lipophilic fluorescent dye PKH26, which integrates into the EV membrane, we tracked EV internalization over time. Our results showed a time-dependent uptake pattern, with HUVECs exhibiting significant internalization within the first hour, peaking between 12 and 24 hours. Notably, PD-EVs uptake plateaued after 12 hours, while DP-EVs and GF-EVs continued to accumulate up to 24 hours. We also observed the localization of internalized EVs toward the nucleus, likely transported via microtubules using motor proteins such as dyneins. This perinuclear accumulation might enhance intracellular signaling or gene regulation, as EVs often deliver cargo essential for cellular processes near the nucleus [53, 54]. However, PKH26 dye transfer from EVs to recipient cell membranes could artifactually inflate uptake estimates, as previously reported [55]. Additionally, the degradation of EVs within lysosomes could reduce fluorescence over time, complicating long-term tracking [48, 56]. Feng et al. demonstrated that EV uptake by phagocytic cells increased significantly over time, reaching a plateau at 12 hours [57]. Similarly, our semi-quantitative analysis suggests that 12 hours may be a critical time point for observing EV internalization by endothelial cells. Prior to this time, EV uptake showed a consistent upward trend, while after 12 hours, the patterns became more variable.

Our findings align with existing evidence that EVs promote angiogenesis by delivering bioactive molecules that regulate endothelial cell proliferation and migration [58]. While stem cell-derived EVs are often highlighted for their angiogenic potential [59], our results revealed distinct differences between the three EV sources. Under controlled conditions, DP-EVs and GF-EVs significantly enhanced vessel length and volume, while PD-EVs were less effective. Importantly, no differences in vessel diameter were observed, suggesting that these EVs primarily promote vascular network expansion rather than altering vessel architecture. This is particularly interesting in tissue regeneration, where robust neovascularization is key to successful therapeutic outcomes. Despite reports of PD-EVs’ pro-angiogenic effects [60, 61], our data suggest stem cell-derived PD-EVs were no better than EVs secreted by immortalized GF in promoting angiogenesis. A more interesting finding is that EVs derived from two different stem cells from the same donors, DP-EVs have a more pronounced pro-angiogenic effect than PD-EVs. This finding is novel, as previous studies have not reported similar observations. It challenges the common notion that stem cell-derived EVs consistently outperform fibroblast-derived EVs in promoting angiogenesis. Considering the challenges of stem cell procurement and ethical concerns, our results highlight the potential of using immortalized fibroblasts as a practical and scalable alternative for EV research and clinical applications.

Different EV subpopulations from different cell type can carry various types and amounts of proteins, reflecting their distinct functional roles. To investigate this, we conducted a proteomic analysis of EVs from three cell sources. The enrichment of “cell adhesion molecule binding” in the Gene Ontology (GO) Molecular Function (MF) category suggests that EVs transport adhesion-related proteins. These proteins may facilitate interactions with recipient cells by binding to surface molecules, promoting EV uptake, and mediating signaling processes critical for tissue repair, immune regulation, and tumor progression [62]. In the GO Cellular Component (CC) category, enriched terms such as “cell-substrate junction,” “extracellular vesicle,” and “extracellular matrix” point to EVs’ roles in mediating cell attachment and interactions with the extracellular matrix. These interactions likely support EV anchoring on recipient cells and regulate cell-matrix communication, which is essential for processes like tissue repair, immune responses, and cancer progression [63]. Similarly, the enrichment of “wound healing” and “angiogenesis” in the GO Biological Process (BP) category highlights EVs’ involvement in promoting tissue repair and blood vessel formation. By delivering bioactive molecules that facilitate cell migration, proliferation, and remodelling, EVs show significant potential for regenerative medicine. Pathway analysis revealed the prominence of “hemostasis” and “innate immune system” pathways in the Reactome database, suggesting that EVs may influence coagulation and immune signaling. Proteins involved in hemostasis could contribute to blood clotting, while components of the innate immune system may enhance cellular responses during injury or stress [53]. Additionally, the VEGFA-VEGFR2 pathway, critical for angiogenesis, indicates that EVs containing these signaling molecules can promote endothelial cell proliferation, migration, and survival, with applications in therapeutic angiogenesis and tissue regeneration [64].

The KEGG pathway enrichment for “proteoglycans in cancer” reveals that EVs may carry proteoglycans essential in tumor microenvironments. These molecules influence cell adhesion, proliferation, migration, and angiogenesis. By facilitating interactions with target cells, EV-associated proteoglycans may contribute to tumor progression through the activation of oncogenic signaling pathways, such as PI3K-Akt and MAPK [65]. These findings underscore the diverse functional roles of EVs and their potential in therapeutic and regenerative applications.

To further analyze the cargo carried by EVs, we performed miRNA sequence analysis. We found that the EV miRNAs showed interesting enrichment in signaling pathways such as “Transcriptional misregulation in cancer,” “MicroRNAs in cancer pathway,” “Signaling pathways regulating pluripotency of stem cells,” “PI3K-Akt” and “TGF-beta signaling pathway” performed by KEGG enrichment analysis. Enrichment of “Transcriptional misregulation in cancer” in EV miRNA sequencing suggests that these EV miRNAs may play a regulatory role in cancer-related gene expression. Such misregulation is critical because EVs can transport miRNAs to recipient cells, influencing oncogenic pathways, cell proliferation, and metastasis. The specific involvement of miRNAs in this pathway highlights their potential impact on tumor progression through transcriptional changes in genes linked to cancer initiation, growth, and drug resistance [66]. Enriching the “MicroRNAs in cancer pathway” highlights the potential of these EVs in cancer regulation and cellular communication in oncogenesis. EVs can carry oncogenic or tumor-suppressive miRNAs, affecting target cells’ signaling pathways, such as those regulating cell proliferation, apoptosis, and metastasis. This enrichment suggests that these EV miRNAs might have crucial roles in tumorigenesis by modulating cancer-related gene expression, potentially offering diagnostic or therapeutic targets in cancer treatment [67, 68]. Enrichment of the “signaling pathways regulating pluripotency of stem cells” points to a potential role for EVs in modulating cell differentiation and maintaining the self-renewal capacity of stem cells. EVs derived from stem cells can carry miRNAs and other molecules involved in pluripotency pathways, influencing the gene expression profiles of recipient cells, and potentially aiding in tissue regeneration and repair processes. This finding suggests a role for EVs in stem cell-based therapies by supporting cellular reprogramming and tissue-specific regeneration. The enrichment of the “PI3K-Akt” and “TGF-beta” signaling pathways in EV miRNA sequencing reflects the potential roles of EVs in modulating cell survival, growth, and differentiation in recipient cells. The PI3K-Akt pathway is known for its role in promoting cell proliferation and survival, commonly implicated in cancer biology and immune responses. Meanwhile, the TGF-beta pathway influences cellular processes like fibrosis, immune regulation, and tissue repair. Together, these pathways suggest that EVs could influence multiple physiological and pathological processes by regulating cell survival, immune responses, and tissue homeostasis [69, 70]. This explains the enhanced vasculogenesis and pro-survival effects observed in the present study.

The PCA of miRNA expression underscores the biological heterogeneity between EV sources, with DP-EVs and PD-EVs clearly segregating, indicating distinct miRNA profiles. In contrast, we found that although the EVs were secreted from DPSC and PDLSC respectively, they still showed obvious clustering that was highly consistent with the donor source. This means that the potential relevance of EVs miRNA expression to biological individual origin is greater than that to cellular origin, which has not been mentioned in previous reports. The looser clustering of GF-derived EVs could reflect the more variable origin of the commercial fibroblast cells. In the present study, only one strain of commercially immortalized GF was used, and we do not know whether the starting cells were derived from only one or different donors, which may have increased the diversity of GF in this study. In addition, the starting cells transfected with the SV40 large T antigen may have led to cellular genomic instability. This could explain why we observed a less pronounced clustering trend, less consistency within groups, and greater biological variability in the hierarchical clustering and PCA analysis of GF-EV miRNA sequencing results. It is important to recognize that the use of commercial GF as a comparison does not fully represent the specificity of true primary fibroblast EVs, which may bias comparisons and reduce the translational relevance of study results. Therefore, our emphasis on donor variation in DP-EV and PD-EV should not be extended to GF-EV. Further studies on GF-EV are needed to clarify this finding.

## Conclusion

This study analysed EVs derived from three oral cell types, including two mesenchymal stem cell lines derived from the same donor. EVs from an immortalized gingival fibroblast cell line were included as a comparator. All three EV types displayed similar surface marker profiles. However, miRNA sequencing and proteomics revealed distinct molecular patterns among the EVs, correlating with differences in their functional performance. Dental pulp stem cell-derived EVs exhibited superior internalization by recipient cells, enhanced cell proliferation, and greater vasculogenesis compared to periodontal ligament stem cell-derived EVs. Interestingly, fibroblast-derived EVs showed comparable performance in functional assays. Principal component analysis of miRNA expression highlighted the biological heterogeneity among EV sources, revealing that donor-specific factors had a greater influence on EV characteristics than cellular origin, a finding not emphasized in earlier studies. These results highlight the complexities of predicting the functional applications of EVs and support the potential of oral stem cell-derived EVs in regenerative medicine.

## Abbreviations

α-MEM: Alpha Minimum Essential Medium
BCA: Bicinchoninic Acid Assay
CM: Conditioned medium
DPSC: Dental pulp stem cell
EGM-2: Endothelial Cell Growth Medium 2
EVs: Extracellular vesicles
FBS: Fetal bovine serum
GO: Gene Ontology
GF: Gingival fibroblast
HUVECs: Human Umbilical Vein Endothelial Cells
IFCM: Imaging Flow Cytometry
KEGG: Kyoto Encyclopedia of Genes and Genomes
MSC: Mesenchymal stromal/stem cell
NTA: Nanoparticle tracking analysis
PBS: Phosphate-buffered saline
PCA: Principal Component Analysis
PDLSC: Periodontal ligament stem cell
TEM: Transmission electron microscopy

## ACKNOWLEDGEMENT

The project was funded by the Deutsche Forschungsgemeinschaft (DFG, German Research Foundation) – project number 516860159.

This work was supported by the Confocal Microscopy Facility and the Genomics Facility, both core facilities of the Interdisciplinary Center for Clinical Research (IZKF) Aachen within the Faculty of Medicine at RWTH Aachen University. Special thanks to Anna Rudzinski, Joseph Chao-Chung Kuo, and Julia Franzen for their invaluable support in sample processing and bioinformatics.

## Competing interests

The authors declare that they have no competing interests.

## Notes

### Competing Interest Statement

The authors have declared no competing interest.

### Summary of Updates

Corrected the scientific name of author in the list of authors.

## References

1. Keshtkar S, Azarpira N, Ghahremani MH. Mesenchymal stem cell-derived extracellular vesicles: novel frontiers in regenerative medicine. Stem Cell Res Ther. 2018;9:63.

2. Zhao AG, Shah K, Cromer B, Sumer H. Mesenchymal stem cell-derived extracellular vesicles and their therapeutic potential. Stem Cells Int. 2020;2020:8825771.

3. Matsuzaka Y, Yashiro R. Therapeutic strategy of mesenchymal-stem-cell-derived extracellular vesicles as regenerative medicine. Int J Mol Sci. 2022;23:6480.

4. Yuan QL, Zhang YG, Chen Q. Mesenchymal stem cell (MSC)-derived extracellular vesicles: potential therapeutics as MSC trophic mediators in regenerative medicine. Anat Rec (Hoboken). 2020;303:1735–1742.

5. Rani S, Ryan AE, Griffin MD, Ritter T. Mesenchymal stem cell-derived extracellular vesicles: toward cell-free therapeutic applications. Mol Ther. 2015;23:812–823.

6. Mantesso A, Sharpe P. Dental stem cells for tooth regeneration and repair. Expert Opin Biol Ther. 2009;9:1143–1154.

7. Iohara K, et al. Side population cells isolated from porcine dental pulp tissue with self-renewal and multipotency for dentinogenesis, chondrogenesis, adipogenesis, and neurogenesis. Stem Cells. 2006;24:2493–2503.

8. Trubiani O, et al. Periodontal ligament stem cells: current knowledge and future perspectives. Stem Cells Dev. 2019;28:995–1003.

9. Imanishi Y, et al. Efficacy of extracellular vesicles from dental pulp stem cells for bone regeneration in rat calvarial bone defects. Inflamm Regen. 2021;41:12.

10. Jin Q, et al. Extracellular vesicles derived from human dental pulp stem cells promote osteogenesis of adipose-derived stem cells via the MAPK pathway. J Tissue Eng. 2020;11:2041731420975569.

11. Zhou H, et al. The proangiogenic effects of extracellular vesicles secreted by dental pulp stem cells derived from periodontally compromised teeth. Stem Cell Res Ther. 2020;11:110.

12. Zhang S, et al. Extracellular vesicles-loaded fibrin gel supports rapid neovascularization for dental pulp regeneration. Int J Mol Sci. 2020;21:4226.

13. Hua S, et al. Periodontal and dental pulp cell-derived small extracellular vesicles: a review of the current status. Nanomaterials (Basel). 2021;11:1858.

14. Liu C, Li Y, Han G. Advances of mesenchymal stem cells released extracellular vesicles in periodontal bone remodeling. DNA Cell Biol. 2022;41:935–950.

15. Diomede F, et al. A novel role in skeletal segment regeneration of extracellular vesicles released from periodontal-ligament stem cells. Int J Nanomedicine. 2018;13:3805–3825.

16. Yan C, et al. Extracellular vesicles from the inflammatory microenvironment regulate the osteogenic and odontogenic differentiation of periodontal ligament stem cells by miR-758-5p/LMBR1/BMP2/4 axis. J Transl Med. 2022;20:208.

17. Xu J, et al. Periodontal ligament stem cell-derived extracellular vesicles enhance tension-induced osteogenesis. ACS Biomater Sci Eng. 2023;9:388–398.

18. Witwer KW, et al. Defining mesenchymal stromal cell (MSC)-derived small extracellular vesicles for therapeutic applications. J Extracell Vesicles. 2019;8:1609206.

19. Ding Z, et al. Understanding molecular characteristics of extracellular vesicles derived from different types of mesenchymal stem cells for therapeutic translation. Extracell Vesicle. 2024;3:100034.

20. Vaka R, et al. Extracellular vesicle microRNA and protein cargo profiling in three clinical-grade stem cell products reveals key functional pathways. Mol Ther Nucleic Acids. 2023;32:80–93.

21. Théry C, Amigorena S, Raposo G, Clayton A. Isolation and characterization of exosomes from cell culture supernatants and biological fluids. Curr Protoc Cell Biol. 2006;Chapter 3:Unit 3.22.

22. Johnstone RM, Adam M, Hammond JR, Orr L, Turbide C. Vesicle formation during reticulocyte maturation. Association of plasma membrane activities with released vesicles (exosomes). J Biol Chem. 1987;262:9412–9420.

23. Ricklefs FL, et al. Imaging flow cytometry facilitates multiparametric characterization of extracellular vesicles in malignant brain tumours. J Extracell Vesicles. 2019;8:1588555.

24. Kreimendahl F, et al. Three-dimensional printing and angiogenesis: tailored agarose-type I collagen blends comprise three-dimensional printability and angiogenesis potential for tissue-engineered substitutes. Tissue Eng Part C Methods. 2017;23:604–615.

25. Ewels PA, et al. The nf-core framework for community-curated bioinformatics pipelines. Nat Biotechnol. 2020;38:276–278.

26. Di Tommaso P, et al. Nextflow enables reproducible computational workflows. Nat Biotechnol. 2017;35:316–319.

27. Merkel D. Docker: lightweight Linux containers for consistent development and deployment. Linux J. 2014;2014(2).

28. Love MI, Huber W, Anders S. Moderated estimation of fold change and dispersion for RNA-seq data with DESeq2. Genome Biol. 2014;15:550.

29. Hughes CS, Moggridge S, Müller T, et al. Single-pot, solid-phase-enhanced sample preparation for proteomics experiments. Nat Protoc. 2019;14:68–85.

30. Licursi V, Conte F, Fiscon G, et al. MIENTURNET: an interactive web tool for microRNA-target enrichment and network-based analysis. BMC Bioinformatics. 2019;20:545.

31. Kolberg L, Raudvere U, Kuzmin I, Adler P, Vilo J, Peterson H. g:Profiler-interoperable web service for functional enrichment analysis and gene identifier mapping (2023 update). Nucleic Acids Res. 2023;51:W207–W212.

32. Kotova AV, et al. Comparative analysis of dental pulp and periodontal stem cells: differences in morphology, functionality, osteogenic differentiation and proteome. Biomedicines. 2021;9:1606.

33. Wang H, et al. Comparative proteomic profiling of human dental pulp stem cells and periodontal ligament stem cells under in vitro osteogenic induction. Arch Oral Biol. 2018;89:9– 19.

34. Ponnaiyan D, Bhat KM, Bhat GS. Comparison of immuno-phenotypes of stem cells from human dental pulp and periodontal ligament. Int J Immunopathol Pharmacol. 2012;25:127–134.

35. Wu T, et al. Comparison of the differentiation of dental pulp stem cells and periodontal ligament stem cells into neuron-like cells and their effects on focal cerebral ischemia. Acta Biochim Biophys Sin. 2020;52:1016–1029.

36. Vasandan AB, et al. Functional differences in mesenchymal stromal cells from human dental pulp and periodontal ligament. J Cell Mol Med. 2014;18:344–354.

37. Ma L, et al. Maintained properties of aged dental pulp stem cells for superior periodontal tissue regeneration. Aging Dis. 2019;10:793–806.

38. Wielento A, Lagosz-Cwik KB, Potempa J, Grabiec AM. The role of gingival fibroblasts in the pathogenesis of periodontitis. J Dent Res. 2023;102:489–496.

39. Häkkinen L, Larjava H, Fournier BP. Distinct phenotype and therapeutic potential of gingival fibroblasts. Cytotherapy. 2014;16:1171–1186.

40. Alfonso García SL, Parada-Sanchez MT, Arboleda Toro D. The phenotype of gingival fibroblasts and their potential use in advanced therapies. Eur J Cell Biol. 2020;99:151123.

41. Naruishi K. Biological roles of fibroblasts in periodontal diseases. Cells. 2022;11:3345.

42. Di Domenico G, Del Vecchio L, Postiglione L, Ramaglia L. Immunophenotypic analysis of human gingival fibroblasts and its regulation by granulocyte-macrophage colony-stimulating factor. Minerva Stomatol. 2003;52:81–91.

43. Phipps RP, Borrello MA, Blieden TM. Fibroblast heterogeneity in the periodontium and other tissues. J Periodontal Res. 1997;32:159–165.

44. Xiong J, et al. Investigation of the cell surface proteome of human periodontal ligament stem cells. Stem Cells Int. 2016;2016:1947157.

45. Torres Crigna A, et al. Inter-laboratory comparison of extracellular vesicle isolation based on ultracentrifugation. Transfus Med Hemother. 2021;48:48–59.

46. Cvjetkovic A, Lötvall J, Lässer C. The influence of rotor type and centrifugation time on the yield and purity of extracellular vesicles. J Extracell Vesicles. 2014;3:23111.

47. Tiwari S, Kumar V, Randhawa S, Verma SK. Preparation and characterization of extracellular vesicles. Am J Reprod Immunol. 2021;85:e13367.

48. Willms E, et al. Extracellular vesicle heterogeneity: Subpopulations, isolation techniques, and diverse functions in cancer progression. Front Immunol. 2018;9:738.

49. Huang Y, et al. Brain tissue-derived extracellular vesicles in Alzheimer’s disease display altered key protein levels including cell type-specific markers. J Alzheimers Dis. 2022;90:1057–1072.

50. Ai Y, et al. Endocytosis blocks the vesicular secretion of exosome marker proteins. Sci Adv. 2024;10:eadi9156.

51. Rosas M, et al. The procoagulant activity of tissue factor expressed on fibroblasts is increased by tissue factor-negative extracellular vesicles. PLoS One. 2020;15:e0240189.

52. Welsh JA, et al. Minimal information for studies of extracellular vesicles (MISEV2023): From basic to advanced approaches. J Extracell Vesicles. 2024;13:e12404.

53. Chae SJ, Kim DW, Igoshin OA, Lee S, Kim JK. Beyond microtubules: The cellular environment at the endoplasmic reticulum attracts proteins to the nucleus, enabling nuclear transport. iScience. 2024;27:109235.

54. Mulcahy LA, Pink RC, Carter DR. Routes and mechanisms of extracellular vesicle uptake. J Extracell Vesicles. 2014;3:24641.

55. Takov K, Yellon DM, Davidson SM. Confounding factors in vesicle uptake studies using fluorescent lipophilic membrane dyes. J Extracell Vesicles. 2017;6:1388731.

56. Tian T, Zhu YL, Hu FH, Wang YY, Huang NP, Xiao ZD. Dynamics of exosome internalization and trafficking. J Cell Physiol. 2013;228:1487–1495.

57. Feng D, et al. Cellular internalization of exosomes occurs through phagocytosis. Traffic. 2010;11:675–687.

58. Ateeq M, Broadwin M, Sellke FW, Abid MR. Extracellular vesicles’ role in angiogenesis and altering angiogenic signaling. Med Sci (Basel). 2024;12:4.

59. Huang M, et al. New insights into the regulatory roles of extracellular vesicles in tumor angiogenesis and their clinical implications. Front Cell Dev Biol. 2021;9:791882.

60. Barile L, Vassalli G. Exosomes: Therapy delivery tools and biomarkers of diseases. Pharmacol Ther. 2017;174:63–78.

61. Zhang Z, et al. PDLSCs regulate angiogenesis of periodontal ligaments via VEGF transferred by exosomes in periodontitis. Int J Med Sci. 2020;17:558–567.

62. Théry C, et al. Minimal information for studies of extracellular vesicles 2018 (MISEV2018): A position statement of the International Society for Extracellular Vesicles and update of the MISEV2014 guidelines. J Extracell Vesicles. 2018;7:1535750.

63. Kalluri R, LeBleu VS. The biology, function, and biomedical applications of exosomes. Science. 2020;367:eaau6977.

64. Muñiz-García A, Wilm B, Murray P, et al. Extracellular vesicles from human umbilical cord-derived MSCs affect vessel formation in vitro and promote VEGFR2-mediated cell survival. Cells. 2022;11:3750.

65. Maas SL, Breakefield XO, Weaver AM. Extracellular vesicles: Unique intercellular delivery vehicles. Trends Cell Biol. 2017;27:172–188.

66. Wang X, et al. Tumor vaccine based on extracellular vesicles derived from γδ-T cells exerts dual antitumor activities. J Extracell Vesicles. 2023;12:e12360.

67. Valadi H, Ekström K, Bossios A, et al. Exosome-mediated transfer of mRNAs and microRNAs is a novel mechanism of genetic exchange between cells. Nat Cell Biol. 2007;9:654–659.

68. Hannafon BN, Ding WQ. Intercellular communication by exosome-derived microRNAs in cancer. Int J Mol Sci. 2013;14:14240–14269.

69. Zhang L, Yu D. Exosomes in cancer development, metastasis, and immunity. Oncogene. 2019;38:4904–4916.

70. Becker A, Thakur BK, Weiss R, Kim HS, Peinado H, Lyden D. Extracellular vesicles in cancer: Cell-to-cell mediators of metastasis. Cancer Cell. 2016;30:836–848.

